# A singular lipid composition supports the progression of plant autophagy

**DOI:** 10.1101/2025.08.25.671700

**Authors:** Josselin Lupette, Thibault Kosuth, Clément Chambaud, Matthieu Buridan, Julie Castets, Valérie Wattelet-Boyer, Inés Toboso Moreno, Céline Yatim, Franziska Dittrich-Domergue, Valérie Gros, Juliette Jouhet, Stéphane Claverol, Su Melser, Manon Genva, Cécile Mirande-Bret, Laetitia Fouillen, Jean-Jacques Bessoule, Frédéric Domergue, Amélie Bernard

## Abstract

Autophagy is an intracellular degradation and recycling pathway essential for cell quality control and plant tolerance to stress. The formation and maturation of autophagosomes, the cargo-packing vesicles, rely on extensive membrane remodeling, yet little is known about the nature, dynamics and functions of lipids in these events. Here, we established a method combing cell fractionation and immuno-isolation in native conditions to purify autophagic membranes from *Arabidopsis thaliana*. By integrating proteomic and lipidomic analyses, we defined their molecular footprint which, coupled to colocalization analyses, revealed potent actors involved in lipid metabolism, membrane trafficking and membrane remodeling, supporting a close interplay between lipid homeostasis and autophagosome biology. Characterization of the phagophore lipid composition showed low sterol and sphingolipid content, a predominance of glycerophospholipids, including phosphoinositides, and a particular enrichment in phosphatidylcholine and phosphatidylglycerol. Comparisons with other plant endomembranes and autophagic compartments from other organisms revealed the singularity of this lipid signature and notably identified phosphatidylglycerol as a plant-specific component of autophagic membranes. Analyses of inducible phosphatidylglycerol-deficient plants showed defects in autophagy activity thereby supporting the functional relevance of the phagophore lipid composition, particularly that of phosphatidylglycerol homeostasis. Together, our findings place lipids as fundamental components of the autophagy molecular landscape and provide a framework to further investigate their contribution to autophagosome biology and functions in plant acclimation to environmental changes.

**Significance Statement:** Autophagy is catabolic pathway critical for eukaryotic life and essential for plant acclimation to stress. It hinges on the remarkable plasticity of a specialized membrane, the phagophore, to orchestrate and support intense membrane remodeling events towards the formation of the autophagic vesicle containing cargo. To resolve their elusive molecular bases, we need to integrate inputs from both components of biological membranes: proteins and lipids, yet information regarding lipids is still very limited. Here, we isolated plant phagophores, established their protein and lipid molecular footprint, revealed its singularity and showed its physiological and functional relevance for autophagy activity. Our work highlights lipids as key regulators of plant autophagy and opens new doors for investigating how membrane dynamics shape cellular stress responses.

## Introduction

Homeostasis relies on the ability of organisms to adapt to environmental fluctuations, including biotic and abiotic stresses. To maintain cellular integrity in such conditions, eukaryotic cells have evolved conserved quality control mechanisms including macroautophagy (hereafter autophagy) (1, 2). This conserved degradation pathway recycles cytosolic components, including proteins and organelles, through their sequestration into specialized vesicles, named autophagosomes, which ultimately deliver cargo to the lytic vacuole (3). Autophagosome formation proceeds through a series of intense and highly coordinated membrane remodeling events. These start with the nucleation of a precursor membrane, the phagophore, which then expands by addition of lipids while engulfing cargo, before sealing into a double membrane autophagosome and fusing with the lytic compartment (1, 2). At the molecular level, autophagy is orchestrated by AuTophaGy-related (ATG) proteins that are sequentially recruited to the phagophore assembly site. Key modules include the ATG1 kinase complex, which controls induction, and the PI3 kinase complex, which mediates phagophore nucleation and recruits downstream effectors (4). Among these, ATG2, ATG9 and ATG18 contribute to membrane expansion, while the ATG conjugation machinery drives lipidation of ATG8 proteins with phosphatidylethanolamine (PE), a central step for membrane dynamics and cargo selection (5, 6). As a membrane-based process, autophagosome biogenesis critically depends on lipids (7), both as structural components and as regulators of membrane dynamics (1, 2). This process requires a substantial lipid supply, which is largely provided by ER– derived lipid transfer mediated by ATG2 and VPS13, coupled to ATG9-dependent lipid scrambling in yeast and mammals (2, 8), but which machinery remains largely uncharacterized in plants (9, 10). In addition to lipid transfer, multiple membrane sources have been proposed to contribute to phagophore formation (2), highlighting the complex origin and potentially unique composition of autophagic membranes. Beyond their abundance, lipids’ nature is also a key determinant of membrane identity, shape and function. In plants, specific lipids, namely phosphatidylinositol-3-phosphate (PI3P) and phosphatidylinositol-4-phosphate (PI4P) have critical roles in autophagosome biogenesis, notably controlling the spatiotemporal dynamics of key ATG proteins at the phagophore (10–12). Besides phosphoinositides, biological membranes display a high diversity of glycerolipids, sphingolipids and sterols, whose physico-chemical properties regulate membrane organization, dynamics, and protein recruitment (13). While these properties are expected to influence all stages of autophagy, from phagophore formation to autophagosome maturation, little is known about the lipid composition of the plant phagophore, its determinants and its functional contribution to autophagosome biology.

Here, to understand how membrane remodeling is instructed and regulated during autophagy, we developed an approach to isolate phagophores and established their protein and lipid footprint. We notably revealed the singularity of the plant phagophore lipid composition and challenged its functional relevance, thereby identifying phosphatidylglycerol (PG) as an unsuspected component of the autophagy pathway. This study supports the critical contribution of lipids to autophagy and provides key molecular actors to further dissect this critical process for plant acclimation to environmental stresses.

## Results

### Isolation of A. thaliana autophagic compartments

To characterize the molecular composition of the phagophore, we developed a method to isolate this compartment, in native conditions, based on an approach previously used to isolate Trans-Golgi-Network (TGN) vesicles (14,15). Briefly, after induction of autophagy in Arabidopsis seedlings (**Fig. S1A**), a membrane fraction (MF) was first collected by cell fractionation and subsequently used as input to immuno-isolate compartments based on their decoration with specific autophagy markers (**Fig. 1A**). We compared two proteins: GFP-ATG8A, which labels both phagophores and autophagosomes but is more accessible for pulldown at the phagophore, being located within autophagosomes (16), and YFP-ATG18A, which associates with early phagophores (5, 17). Electron microscopy confirmed the presence of labeled membrane vesicles specifically in the IP_+_ fraction compared to the negative control (IP_−_, using the same input with uncoated beads; **Fig. S1B**). Western blot analyses showed the enrichment of ATG8A or ATG18A in the immunoisolated fractions compared to the input MF, confirming efficient pulldown (**Fig. 1B–C**). Markers of major cellular compartments (including the ER, mitochondria, nucleus, chloroplasts, plasma membrane and Golgi) were not detected, indicating high purity of the isolates. In contrast, the autophagy cargo receptor NBR1 was enriched in both ATG8A- and ATG18-isolated fractions. Together, these results demonstrate validate our approach to isolate ATG8A- and ATG18A-associated membranes for downstream molecular characterization.

**Figure 1.**
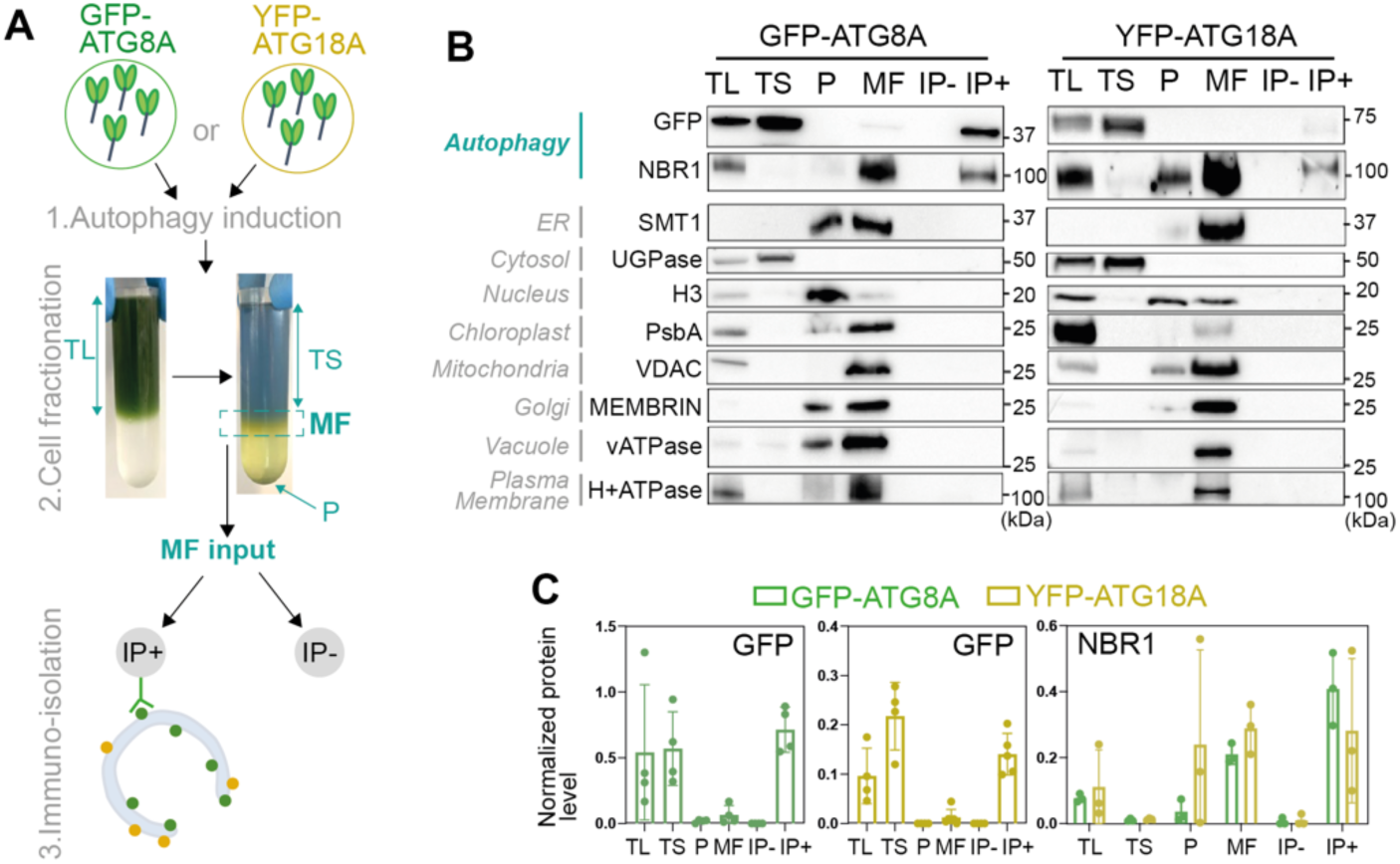
Isolation of autophagic compartments in *Arabidopsis thaliana*. **A**. Isolation was performed with two autophagy markers expressing lines, YFP-ATG18A and GFP-ATG8A. (1) After autophagy induction, (2) seedlings were grinded, and the total lysate (TL) was deposited on a 38 % sucrose cushion and ultracentrifuged overnight to separate the total soluble fraction (TS) and the membrane fraction (MF) from larger cellular compartments (Pellet, P). (3) The membrane fraction (MF) was collected and served as input for the immuno-isolation which was carried out with control magnetic beads (uncoated, IP_−_) or GFP-trap beads (IP_+_). **B**. Western-blot analyses of the different fractions collected during the immuno-isolated procedure. Antibodies raised against different cell compartment markers were used: GFP and NBR1 (autophagy), SMT1 (ER), UGPase (cytosol), H3 (nucleus), PsbA (chloroplast), VDAC (mitochondria), MEMBRIN (Golgi apparatus), vATPase (vacuole) and H^+^ATPase (Plasma Membrane). **C**. Quantification of autophagy markers (GFP and NBR1) in the different cell fractions. Results present the level of each protein normalized by the total protein detected in the stain free in n=4 (anti-GFP) and n=3 (anti-NBR1) independent biological replicates.

### Characterization of the proteomes of ATG-enriched membranes

To characterize the protein composition of ATG18A- and ATG8A-associated membranes, we performed proteomic analyses of immuno-isolated fractions (IP_+_), their controls (IP_−_), and input membrane fractions (MF, **Fig. 2A-D**). After filtering out non-specific proteins, we identified 904 and 729 proteins significantly enriched in ATG18A-(**Dataset S1**) and ATG8A-(**Dataset S2**) isolates compared to MF, respectively. Both datasets contained core autophagy components, further supporting the identity of the isolated membranes as phagophores. Functional classification revealed enrichment in proteins related to lipid metabolism and membrane trafficking. These include enzymes involved in phosphoinositide metabolism (SAC9, PI4Kα1), fatty acid synthesis and lipid remodeling, as well as components of major trafficking machineries such as ESCRT, retromer, exocyst and SNARE complexes. Comparison of both datasets identified a shared core of 233 proteins, including protein actors involved in vesicular trafficking, and intracellular transport (GARP complex at the Golgi and MAG2 complex at the ER) and lipid metabolism (SAC9, FRA3) suggesting conserved molecular features across autophagic compartments (**Fig. 2C-D, Dataset S3**). Transient expression of selected candidates from various functional nodes in *Nicotiana benthamiana* showed partial to strong colocalization with ATG8E for 14 out of 25 tested proteins (**Fig. 3; Fig. S2**). This supports their association with autophagic compartments and the robustness of our dataset which, by defining the proteomic landscape of plant phagophores, provides potent candidates involved in membrane dynamics and lipid metabolism during autophagy.

**Figure 2.**
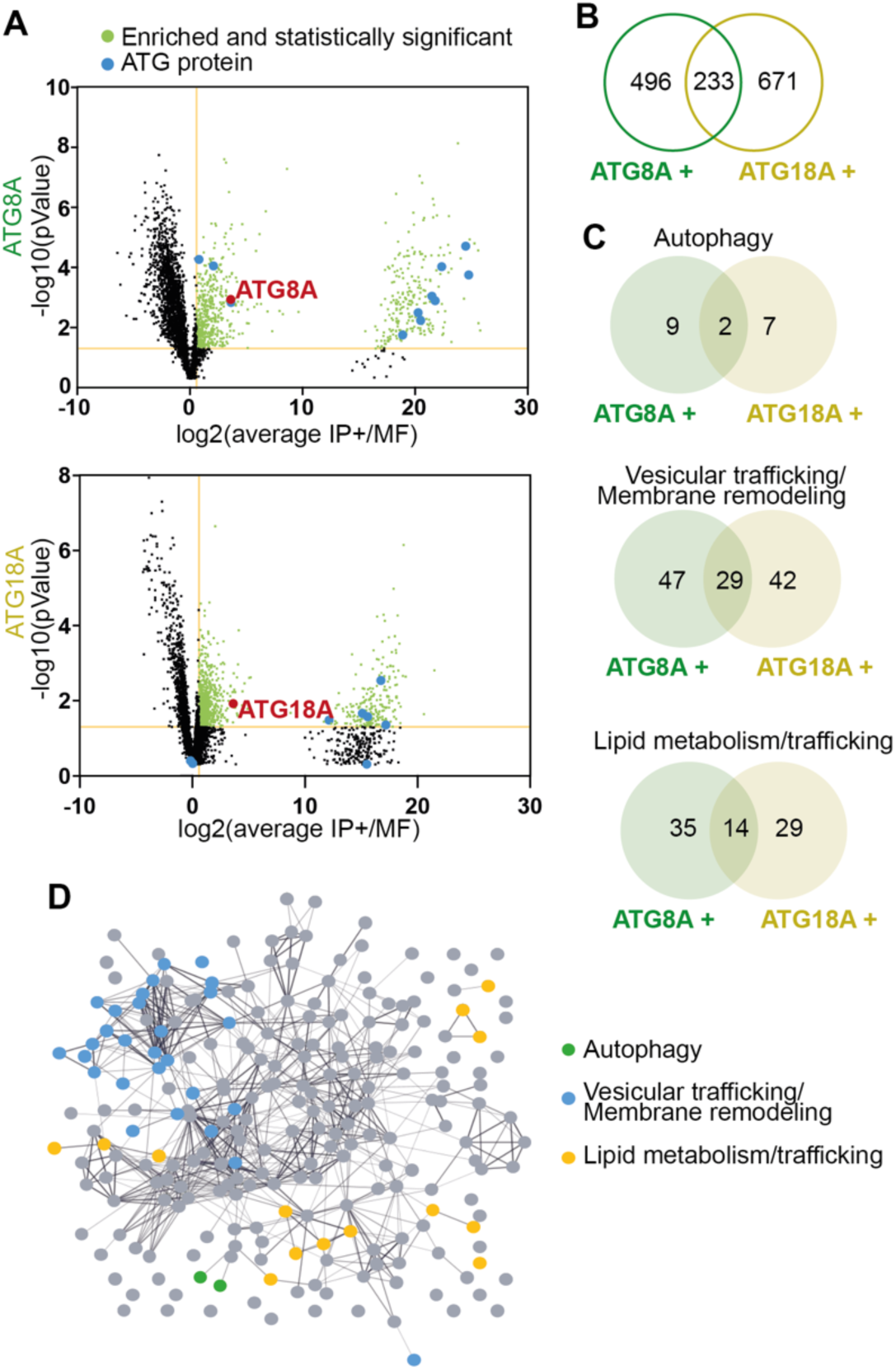
Proteomic analyses of isolated ATG-compartments. **A**. Volcano plots of proteins found in the ATG8A- and ATG18A-immuno-isolated compartments presenting the average protein enrichment in the IP+ fraction compared to the input MF, expressed as log2 (relative abundance in IP_+_/ relative abundance in MF), on the *x* axis plotted against the −log10 (p-value) on the *y* axis. Yellow lines mark the thresholds (log2(IP_+_/MF)>0.585; −log10(p-value)>1.301) used to select proteins significantly enriched in the IP fraction, that are labeled in green. Autophagy-related proteins (ATG) are indicated in blue and prey proteins (ATG8A and ATG18A) are colored in red. Proteins not found significantly enriched in the IP fraction are represented as black dots. **B-C**. Venn diagrams indicating the number of common or specific proteins enriched in ATG18A-(ATG18+) and ATG8A-(ATG8A+) isolated compartments (B) and specifically those involved in autophagy, vesicular trafficking and membrane remodeling or lipid metabolism/trafficking (C). Protein lists are shown in **Datasets S1-S3. D**. Functional interactome of the 233 proteins, listed in **Dataset S3**, found enriched in both ATG18A- and ATG8A-isolated compartments. Proteins involved in autophagy are shown in green, those involved in vesicular trafficking and membrane remodeling are in blue, and those implicated in lipid metabolism are in yellow. Other proteins are shown in gray. The thickness of the line indicates the percentage of confidence of interaction between two proteins.

**Figure 3.**
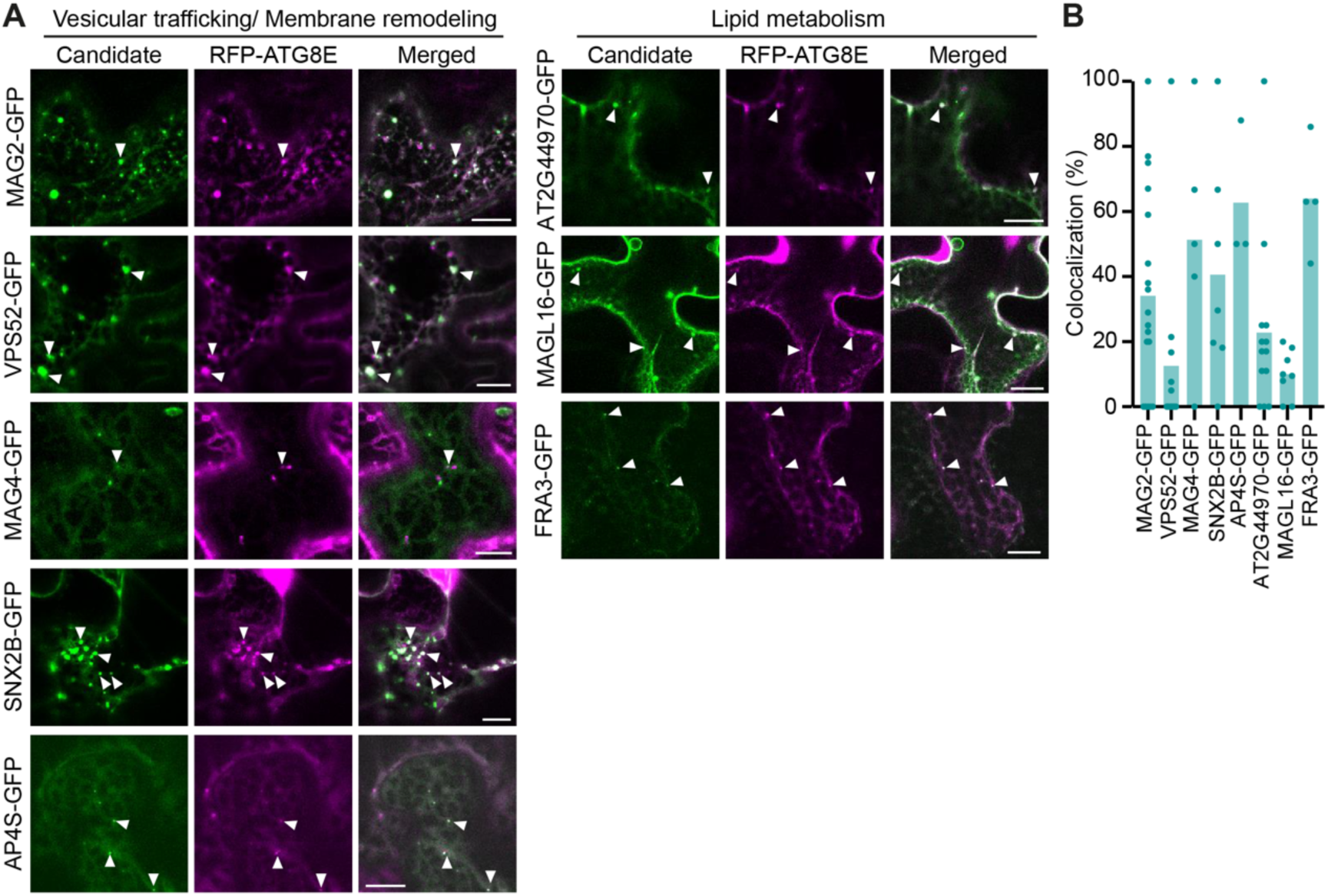
Colocalization analyses of protein candidates and RFP-ATG8E in leaves of *Nicotiana benthamiana*. **A**. Representative confocal images of the subcellular localization of protein candidates-GFP (involved in vesicular trafficking/ membrane remodeling or lipid metabolism) in regard to that of the autophagy marker RFP-ATG8E in *Nicotiana benthamiana* leaves. Colocalization are indicated with white arrows. Scale bar: 10 µm. **B**. Quantification of images of (**A**). Results present the percentage of RFP-ATG8E signal found on candidate puncta. Results are presented as histograms with the mean and individual data of n=4-16 replicates in 1 to 3 independent biological experiment.

### Lipid composition of autophagic membranes

Global fatty acyl chain and sterol profiling of ATG8A- and ATG18A-isolated membranes (**Fig. 4**) showed highly similar profiles, which differed markedly from input MF, with a strong enrichment in fatty acids (FAs) and a concomitant depletion in sterols and α-hydroxy fatty acids (h-FAs) (**Fig. 4A-B**). Analyses of the molecular composition of fatty acyl chains showed a larger proportion of unsaturated FAs compared to MF (C18:2+C18:3, although not statistically significant), and revealed the notable absence of C16:3 species that are characteristic of plastidial lipids (18) (**Fig. 4C**). This supports the specificity of the isolated compartments and indicates that chloroplast-derived lipids do not contribute to autophagic membranes in these conditions (**Fig. 4C**). Specific to sphingolipids (19), the abundance of h-FAs (<1%) indicates that sphingolipids are minor components of the phagophore and targeted analyses showed no significant qualitative differences compared to MF (**Fig. S3**). These results suggest that plant autophagic membranes are predominantly composed of glycerolipids (~95% of total lipids, as reflected by the FA content), with minimal contributions from sterols and sphingolipids.

**Figure 4.**
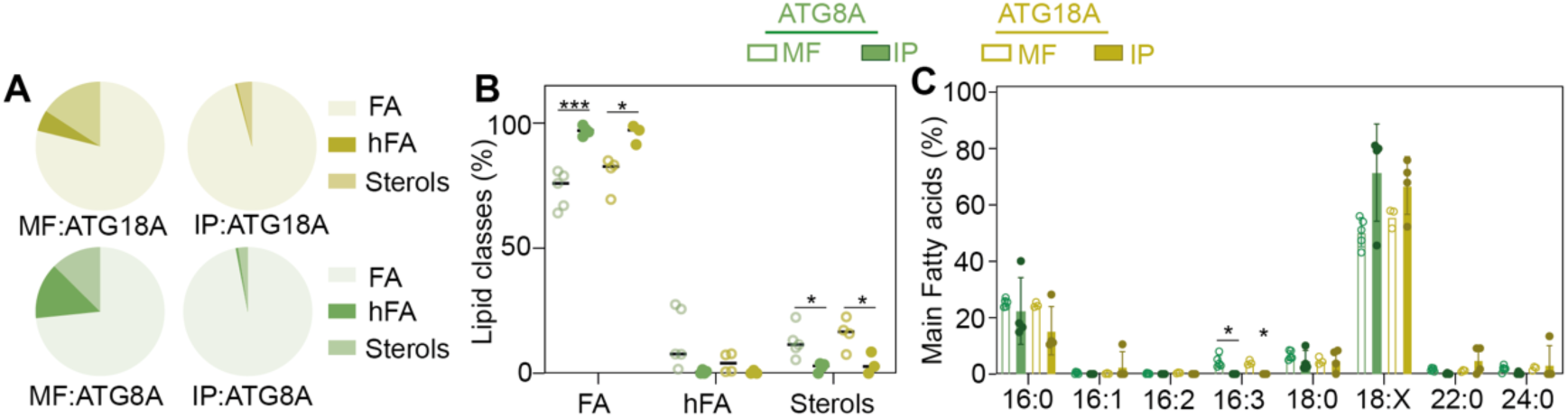
Fatty acyl chain and sterol profiles of ATG-immuno-isolated compartments compared to their respective membrane fraction input. Fatty Acid Methyl Esters (FAMEs) compositions were determined by GC-MS after transmethylation of the membrane fractions (MF) and of the immuno-isolated compartments (IP). **A-B**. Relative distribution of FAMEs (fatty acids, FA and hydroxy fatty acids, hFA) and sterols within the MF and IP fractions in YFP-ATG18A (yellow) and GFP-ATG8A (green) lines represented as pie chart (**A**) and with individual values (**B**). Results present each class of molecules as percentage of total lipids. **C**. Relative distribution of the main fatty acid found in MFs and isolated compartments (IP) in GFP-ATG8A (green) and YFP-ATG18A (yellow). 18:X represents the sum of C18:2+C18:3. Results present each class of fatty acids as percentage of total fatty acids. n=3-5 independent biological experiment. Student t-test was performed to assess statistical differences between MF and IP fractions. P-values are as follows: p-value>0.05 (non-significant, ns), *p<0.05, **p<0.01 and ***p<0.001.

### Glycerolipidome of A. thaliana autophagic membranes

Consistent with the lack of C16:3 species, comprehensive glycerolipid profiling by LC–MS/MS did not detect plastidial lipids, (MGDG, DGDG, SQDG), in ATG8A- or ATG18A-isolated membranes; similarly, mitochondrial (DPG) and storage lipids (TAG) were absent, supporting the specificity of the phagophore. Instead, we found that autophagic membranes are mainly composed of phosphatidylcholine (PC) and phosphatidylglycerol (PG), followed by phosphatidylinositol (PI) and phosphatidic acid (PA), and lower levels of phosphatidylserine (PS) and phosphatidylethanolamine (PE). Notably, the phagophore composition showed significant differences in specific lipid classes (enrichment in PC and PG and a depletion in PE; **Fig. 5A-B**) while distributions of lipid molecular species were found largely similar to that of the input MF (**Fig. 5C-D**). Given the known importance of phosphoinositides in autophagy (2), these were further analyzed using a dedicated LC-MS^2^ method (20). Monophosphate phosphoinositides (PIP), corresponding to PI3P and/or PI4P, were detected in immuno-isolated fractions, whereas biphosphate phosphoinositides (PIP_2_, including PI(3,5)P_2_ and PI(4,5)P_2_) species were not (**Fig. 5C**), consistent with their known restriction to the plasma membrane in plants (11). In sum, our results unravel the plant phagophore glycerolipidome, characterized by the presence of specific phosphoinositides and the enrichment in PC and PG.

**Figure 5.**
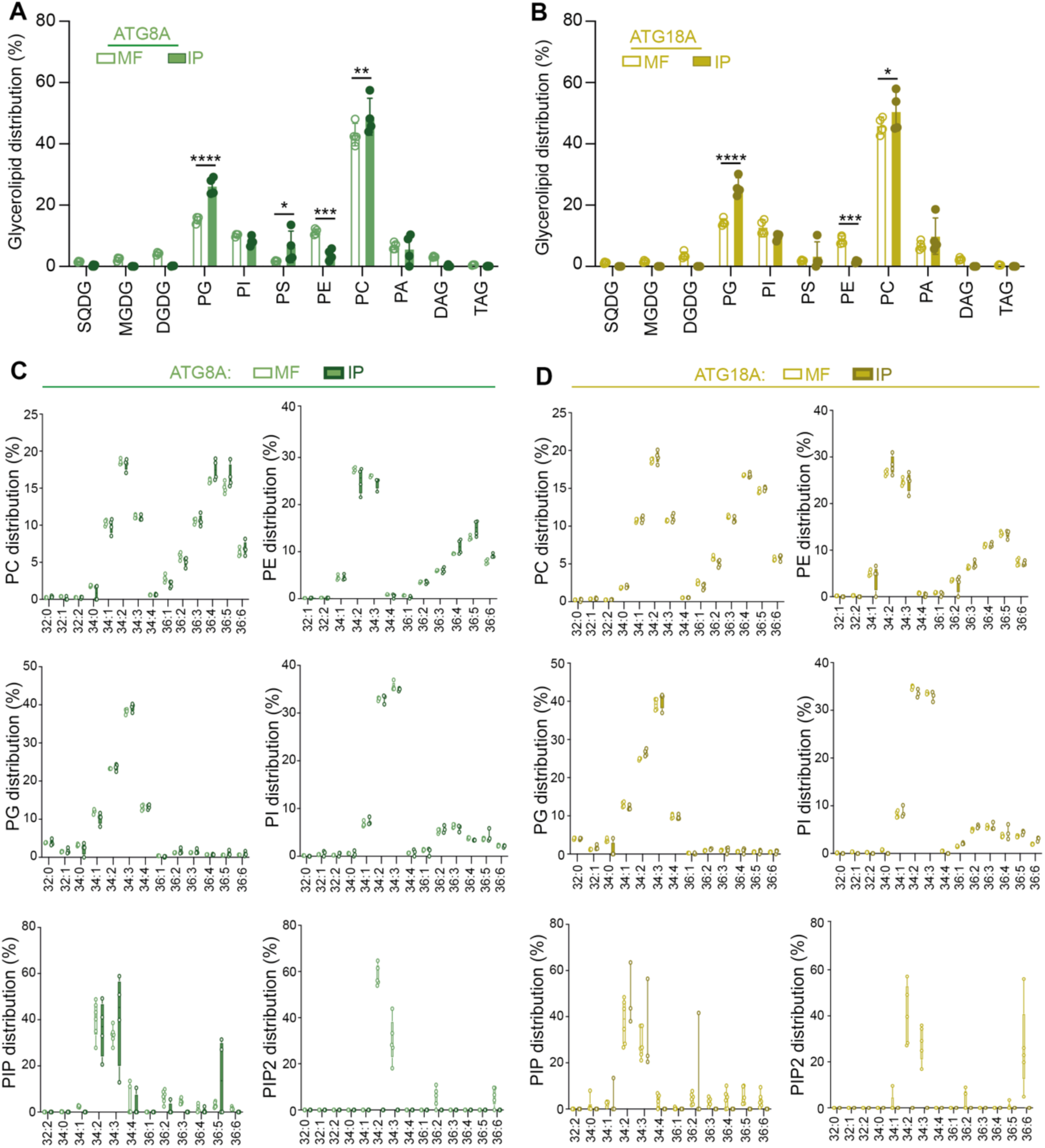
Glycerolipidome of ATG-immuno-isolated compartments compared to the membrane fraction. Glycerolipidome of membrane fraction (MF) and autophagic compartment (IP+) fractions determined by HPLC-MS/MS for **A**. GFP-ATG8A (green) and **B**. YFP-ATG18A (yellow) lines. n=3-4 independent biological experiment. ANOVA analyses showed statistical differences between MF and IP fractions. P-values are as follows: p-value>0.05, non-significant, *p<0.05, **p<0.01, ***p<0.005 and ****p<0.001. Molecular species distribution of PC, PE, PG, PI, PIP and PIP_2_ of the MF and IP fractions for GFP-ATG8A (green, **C**) and YFP-ATG18A (yellow, **D**) lines. SQDG, sulfoquinovosyldiacylglycerol; MGDG, monogalactosyldiacylglycerol; DGDG, digalactosyldiacylglycerol; PG, phosphatidylglycerol; PI, phosphatidylinositol; PS, phosphatidylserine; PE, phosphatidylethanolamine; PC, phosphatidylcholine; PA, phosphatidic acid; DAG, diacylglycerol; TAG, triacylglycerol; PIP, monophosphate phosphoinositides and PIP_2_, diphosphate phosphoinositides.

### The plant phagophore, a singular lipid signature

Each and every cell compartment is defined by a particular lipid composition which shapes its identity, architecture and functions. To question the phagophore specificity and highlight potentially relevant features, we compared its composition to that of other membrane compartments (plasma membrane, ER, Golgi, TGN, tonoplast, plastids). First, we found that autophagic membranes are highly divergent from that of the plasma membrane or tonoplast were sphingolipids and sterols accumulate substantially (19, 21–22) compared to the minor proportion of these lipids in ATG8A- or ATG18A-isolates (**Fig. 4A-B**). In that regard, they also differ from the sphingolipid-enriched TGN (14, 15). Second, the phagophore stands out from plastids or mitochondria, with the absence of *bone fide* lipid markers of these compartments (MGDG, DGDG, SQDG for plastids, 23–24; DPG for mitochondria, 25–27). Instead, the phagophore composition appears closer to that of the ER, which aligns with the major importance of this organelle for autophagosome biogenesis (2, 4, 6), but with a striking difference in the accumulation of PG (**Fig. 5A-B, Fig. 6A**). Reaching approximately 25–26 mol% in both ATG18A- and ATG8A-isolated membranes (**Fig. 5**), PG levels are substantially higher than that reported for other plant membranes, where PG typically represents only a minor fraction (**Fig. 6A**). Further comparison with non-photosynthetic systems indicates both conserved and divergent features of autophagic membranes (**Fig. 6B**, 28–30). While enrichment in unsaturated FAs appears to be a common characteristic across species, the relative abundance of phospholipids varies considerably. PC, PE, PI and PS are universally present but in different proportions. However, PG emerges here as a distinctive feature of plant autophagic membranes, being largely absent from autophagosomes in other model systems.

**Figure 6.**
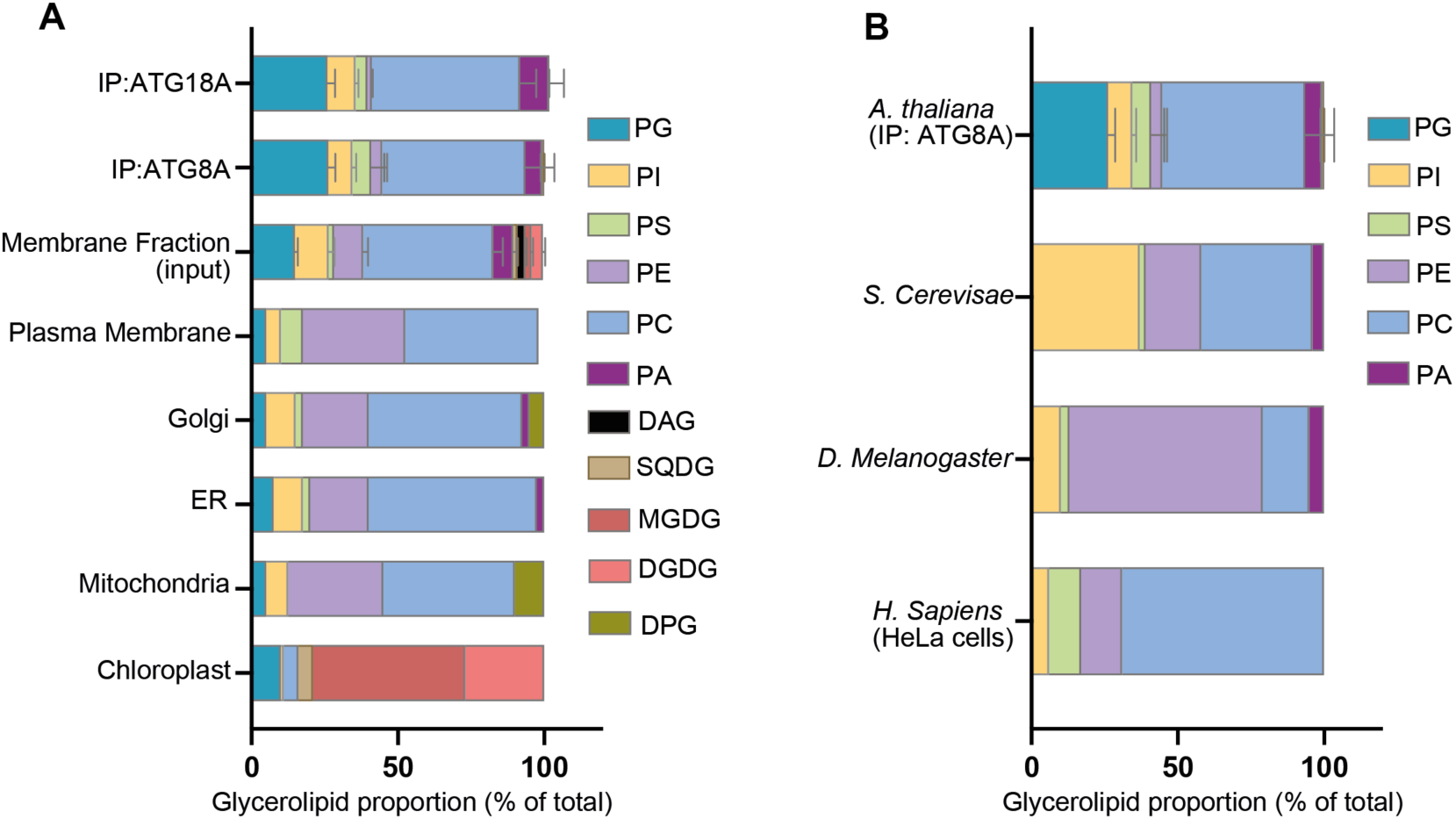
The plant phagophore shows a singular lipid signature. **A**. Comparing the glycerolipidome of autophagic membranes with that of several plant membrane compartments shows the particularity of its composition. Data present the relative distribution of each glycerolipid as percentage of total glycerolipids using results from this study (IP:ATG18A, IP:ATG8A, MF, see **Fig. 5A,B**) and that from previous publications for Golgi & ER (from barley, *Hordeum vulgare*, 21), Plasma Membrane (from mung bean, *Vigna radiata*, 22), Plastid (from spinach, *Spinacia oleracea*, 23) and Mitochondria (from cauliflower, *Brassica oleracea*, 25) **B**. Comparison of the glycerolipid composition of *A. thaliana* autophagic membranes with that from non-photosynthetic organisms. Data present the relative distribution of each glycerolipid as percentage of total glycerolipids using results from this study (IP:ATG8A, see **Fig. 5A**) and that from previous publications for autophagic compartments purified from yeast *(S. cerevisae*, 28), Drosophila (*D. melanogaster*, 29) and HeLa cells (*H. sapiens*, 30). SQDG, sulfoquinovosyldiacylglycerol; MGDG, monogalactosyldiacylglycerol; DGDG, digalactosyldiacylglycerol; PG, phosphatidylglycerol; PI, phosphatidylinositol; PS, phosphatidylserine; PE, phosphatidylethanolamine; PC, phosphatidylcholine; PA, phosphatidic acid; DAG, diacylglycerol, DPG, diphosphatidyldiacylglycerol.

### Phosphatidylglycerol is required for efficient autophagy in leaves

Next, we aimed at challenging the physiological and functional relevance of the phagophore lipid composition. Particularly, given the enrichment and specificity of PG in plant phagophores, we assessed the impact of downregulating PG synthesis on autophagy activity. In Arabidopsis, the *pgps1* mutant is severely impaired in PG biosynthesis and shows significant alterations in chloroplast function and plant development (31), whereas the *pgps1 pgps2* double mutant is embryo-lethal (32). Because the physiological consequences of these mutations are too severe and pleiotropic, these lines are unsuitable for direct and specific functional analysis of the role of PG in autophagy. To disrupt all pools of PG while overcoming lethality and limiting pleiotropic effects we generated an inducible amiRNA line (*ami:pgps1,2*) downregulating both *PGPS1* and *PGPS2* (**Fig. S4A, B**) and showing a large reduction in PG levels (**Fig. 7A**). The *ami:pgps1,2* line showed mild defects in development compared to the WT in rich conditions (see +NC conditions **Fig. 7B-C, Fig. S4C-D**). This phenotype was exacerbated by prolonged nutrient starvation (-NC, lacking nitrogen and carbon), particularly in photosynthetic tissues, as shown by recovery analyses (**Fig. 7B-C, Fig. S4C-D**). Quantification of autophagy flux using the GFP-ATG8 processing assay showed no significant difference in roots (**Fig. 7D**), but a marked 50% reduction in *ami:pgps1,2* compared to WT in leaves (**Fig. 7E**). Consistently, the autophagy-receptor NBR1 was found over accumulating in *ami:pgps1,2*, suggesting defects in its degradation by autophagy (**Fig. S4E**). Altogether, these results show that downregulating the content of PG leads to a partial block in autophagy degradation specifically in leaves and therefore suggest that PG is an important component of autophagy activity in Arabidopsis.

**Figure 7.**
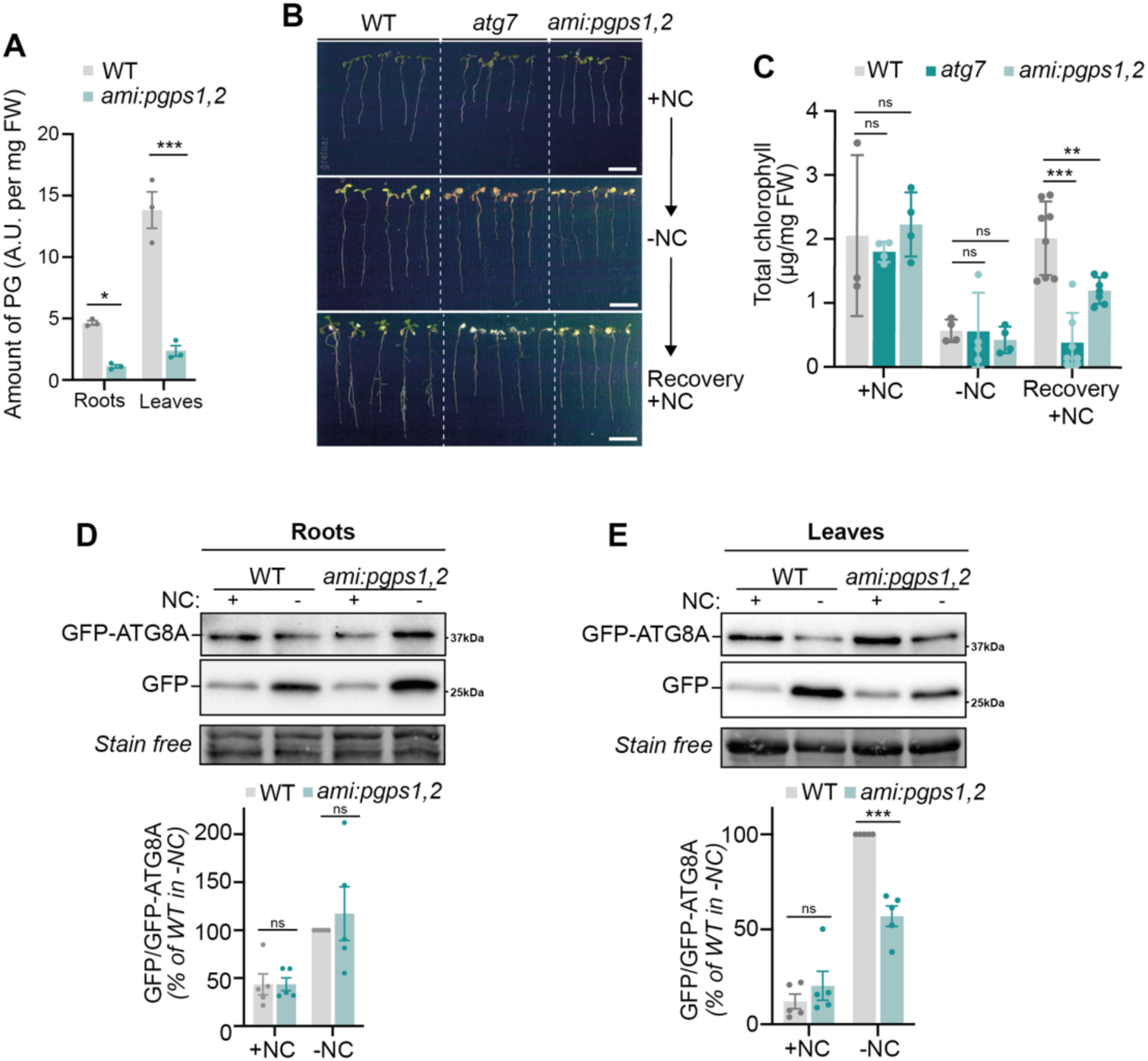
Downregulating *PGPS1* and *PGPS2* compromises autophagy in leaves. **A**. Downregulating *PGPS1* and *PGPS2* leads to a decrease in PG content compared to WT. Quantification of the amount of PG in pUB10::GFP-ATG8A (WT) and *ami:pgps1,2*_pUB10::GFP-ATG8A (*ami:pgps1,2*). Roots and leaves of 7-day-old seedlings grown on MS with 10 μM β-estradiol were dissected; total phospholipids were extracted and analyzed by HPLC-MS/MS. Results present individual values and mean ± SEM of n=3 independent biological replicates. **B-C**. Plants with a lower amount of PG show phenotypical differences compared to WT after prolonged starvation. WT, *ami:pgps1,2* and *atg7* seedlings grown in rich conditions (+NC) were transferred to nutrient deprived (-NC) solid medium for 10 days in darkness (-NC) and then transferred back to +NC solid medium for 7 days for recovery. All media were supplemented with 10 μM β-estradiol. Scale bar: 1 cm. Representative images (**B**) are quantified in (**C**). Total chlorophyll was extracted and quantified in each condition. Results are presented as the mean ± SEM with values of distinct biological replicates (n≧3) within 2 independent experiments. Statistical differences were assessed using two-way ANOVA. **D-E**. GFP-ATG8 processing assays show significant reduction in autophagy flux in leaves of *ami:pgps1,2* compared to WT (**E**); in contrast, autophagy is not affected in the roots (**D**). Seedlings grown on rich medium were transferred to either nutrient rich liquid medium (+NC) or nutrient-starved liquid medium (-nitrogen and carbon in the dark; −NC) for 6 h; media were all supplemented with 10 μM β-estradiol. Total proteins from leaves and roots, dissected after treatments, were extracted and analyzed by immunoblotting with anti-GFP antibodies. Stain free images were used as loading control. Representative images are provided in the upper panels. Quantification of the ratio of GFP/GFP-ATG8A reflective of the rate of autophagy flux is shown in the bottom panels, relative to that of control plants (WT −NC) which was set to 100% in each experiment. Results are presented as the mean ± SEM with values of all replicates (n=5 independent biological replicates).

## Discussion

While autophagy is a membrane-based process, quantitative, qualitative and functional information regarding the lipid composition of the phagophore had remained scarce. Here, we establish a method to isolate plant phagophores (**Fig. 1**), show its potential to uncover novel molecular actors of the autophagy pathway (**Fig. 2–3, 7**), reveal specificities in its lipid composition (**Fig. 5–6**) and provide evidence that support its functional relevance (**Fig. 7**).

Our work notably shows that the plant phagophore is largely composed of glycerolipids with unsaturated FAs, with a very minor contribution of sphingolipids and sterols (**Fig. 4**). Although these are relatively low abundant, their functional contribution might still be relevant (2) and warrant dedicated analyses in future studies. Similarly, the low content of PE detected in phagophores (**Fig. 5A-B**) does not reflect a reduced functional role in ATG8 lipidation but rather arises from a technical limitation of LC-MS/MS–based analyses, which only quantifies the pool of free PE. This highlights an important conceptual point: bulk analyses likely not fully represent the functionally active lipid species at sites of autophagosome formation. In this context, PE should be viewed not only as a membrane constituent but also as a dynamic substrate whose local consumption is tightly coupled to ATG8 lipidation cycles. This supports the idea that autophagic membranes are highly dynamic lipid platforms, in which lipid turnover/usage in addition to steady-state abundance may be most relevant for function.

Irrespective of the marker used for immuno-isolation, we found a similar proportion between cylindrical glycerolipids (85%: PC, PI and PG) and conical lipids (15%: PE, PA and PS; **Fig. 5**) in the phagophore. Future studies including molecular modeling and *in vitro* reconstitution should aim at deciphering whether this balance is a structural requirement for autophagosome formation, where large-scale membrane expansion must be coordinated with the generation and maintenance of highly curved regions at the phagophore rim. Work in yeast showed that a complete depletion in PC, the major component of autophagosome (28), does not alter phagophore expansion but rather disrupts the completion of the vesicle (33). In that context, by resolving the nature of lipids composing plant phagophores, our study now paves the way to examine their lateral heterogeneity and to assess whether this supports dedicated functional and/or ultrastructural subdomains to promote its shaping and remodeling to an autophagosome. Beyond structural implications, the regulatory roles of the phagophore lipidome should also be considered. We found that autophagic membranes are enriched in anionic lipids, including phosphoinositides and PG, which contribute to the membrane association of selected proteins through electrostatic interactions (2). Lipid–protein interactions are already recognized as critical determinants of autophagy, exemplified by the binding of multiple autophagy effectors to PI3P (2, 4). Such a lipid environment may provide a platform for the dynamic assembly of the autophagy machinery and the regulation of membrane remodeling events. Among these lipids, PG emerges as a notable candidate which functional relevance is discussed below.

Comparing plant phagophores to other endomembranes or to autophagic compartments from other model organisms identified its singularity and specificities. These notably supports the predominance of the ER as a source of lipids for the expansion of the phagophore (2, 4, 6), shows the versatility of the lipid composition of autophagic membranes across species (**Fig. 6**), and identifies PG as a plant-specific component of the phagophore. Particularly, we show that PG accumulates at the phagophore and that its depletion causes defects in autophagy activity and tolerance to nitrogen and carbon starvation, specifically in leaves (**Fig. 7**). This specificity may reflect differential composition between phagophores from leaves and roots. Accounting to the relative mass of each organ, phagophores from leaves are likely predominant in the pool of autophagic membranes isolated from whole seedlings. As such, the observed enrichment in PG (**Fig. 5**), may be specific to phagophores from photosynthetic tissues, hence the absence of root autophagy phenotype. Out of the scope of this study, the analytical pipeline established here should be instrumental to explore whether the phagophore composition varies depending on organ, tissue, environmental conditions and importantly, populations of autophagosomes and types of selective autophagy which may affect the lipid/membrane source required the formation and functions of these compartments.

Irrespective of such specificities, our data support the physiological relevance of the phagophore lipid composition and particularly point to PG as an important component of autophagy in leaves both quantitatively and functionally, even though we acknowledge that functions of PG outside of the phagophore may also indirectly affect the autophagy pathway. At this time, four speculative functions could be attributed to PG in autophagosome biology. (*i*) It may be quantitatively relevant, providing building blocks for the expansion of the phagophore; (*ii*) as an anionic lipid it may contribute to the phagophore electrostatic landscape to support dynamic protein association during autophagosome formation/maturation; (*iii*) it may support the assembly or activity of protein complexes at the phagophore, similarly to its function as a structural cofactor of the photosystem II (34); (*iv*) it may engage in specific lipid-protein interaction thereby recruiting dedicated actors of the autophagy machinery or participating in the selection of autophagic cargo. In line with the latter, PG has been shown to directly bind the mobile floral activator Florigen FT in Arabidopsis, thereby modulating its intracellular localization and biological activity (35). Whether analogous PG-dependent interactions occur within the autophagy machinery remains unknown and future research should aim at challenging the role of PG in organizing autophagic membranes and/or controlling the spatiotemporal distribution of key autophagy actors in plants.

Besides the lipid composition of the phagophore, our study also provides proteomics analyses revealing the molecular framework that underlies autophagic membrane dynamics. Although ATG18A- and ATG8A-associated membranes displayed highly similar lipid compositions (**Fig. 5**), their proteomes were more distinct (**Fig. 2, Datasets S1-S3**), likely reflecting different specific interactors and/or stages of autophagosome biogenesis. Overall, the presence of well-described autophagy-associated proteins, the partial overlap with previously reported autophagosomal-enriched proteomes (36, **Dataset S3**) and the localization of numerous candidates to autophagic structures (**Fig. 3**) further validate the robustness of the dataset. Beyond confirming known components of the pathway, our data provide fundamental insights into the molecular actors and processes associated with the dynamics of the phagophore during autophagy. A striking feature is the enrichment of proteins involved in lipid metabolism, including phosphoinositides –and particularly PI4P– dynamics, FA synthesis and modification, lipid transfer, and membrane lipid remodeling (**Datasets S1-S3**). Together with the lipidomic data, these observations suggest that autophagic membranes are not merely assembled from pre-existing cellular membranes but undergo active lipid remodeling throughout phagophore growth and maturation. Such a model is consistent with the substantial membrane expansion required during autophagosome biogenesis and highlights lipid homeostasis as a central component of the process. Our dataset also revealed a strong representation of membrane trafficking and tethering actors, supporting extensive interactions between autophagic membranes and the endomembrane system (6, 37–39). Rather than implicating a single membrane source, these findings are consistent with current models in which multiple organelles contribute membrane material and regulatory activities during autophagosome formation (6, 37–39). In this context, lipid metabolism and membrane trafficking likely act in concert to ensure the coordinated delivery, remodeling, and organization of membrane components required for phagophore expansion. Further characterization of the phagophore-enriched proteins identified here should, and actually already has, provided valuable insights to understand lipid dynamics during autophagosome formation and maturation and to dissect the molecular bases of elusive stages of the autophagy pathway. A striking example is our identification of the phospholipase LCAT4, enriched in ATG8-isolated membranes (**Dataset S2**), which functional characterization recently shed light on the previously uncharacterized step of membrane hydrolysis for the disruption and degradation of autophagic bodies in the vacuole (40).

In sum, our work provides a comprehensive molecular characterization of plant autophagic membranes integrating both lipidomic and proteomic information. While proteins have historically dominated the study of autophagy, our results highlight the value of considering the membrane itself as a dynamic and regulated entity. Given the central role of autophagy in plant stress tolerance, identifying how lipid homeostasis contributes to autophagy opens new gates for understanding and managing the effects of environmental changes on plant physiology.

## Materials and Methods

### Plant Materials and growth conditions

Lipidomic and proteomic studies were performed in *Arabidopsis thaliana* of the ecotype Columbia-0 (Col-0) as wild-type. The following transgenic lines in Col-0 were used: 35S::GFP-ATG8A (41), YFP-ATG18A (17). The generation of *ami:pgps1,2* line is detailed in **Supplemental Information**. The lines pUB10::GFP-ATG8A (WT, 42), *ami:pgps1,2* expressing pUB10::GFP-ATG8A (*ami:pgps1,2*) and *atg7-2* expressing pUB10::GFP-ATG8A (*atg7*, 42) were used for recovery assays and immunoblot analyses detailed in **Supplemental Information**. Seeds were sterilized and grown in liquid Murashige and Skoog (MS) medium under standard conditions for 7 days. Autophagy was induced as indicated for the times indicated in the figures, more details are available in **Supplemental Information**.

### GFP-ATG8 assay and NBR1 immunoblot analyses

Plant material was frozen in liquid nitrogen and disrupted using a TissueLyser. Proteins were extracted and denatured at 55 °C for 15 min. Proteins were separated by SDS-PAGE (12%) and transferred to nitrocellulose membranes. Immunodetection was performed using the appropriate antibodies (anti-GFP antibody, Roche, #11814460001, 1/2000; anti-NBR1 antibody, Agrisera, #AS142805, 1/5000) followed by HRP-conjugated secondary antibody (1:10,000). Signal was revealed using ECL and acquired using a ChemiDoc system. More details are provided in **Supplemental Information**.

### Immuno-isolation of autophagic membranes

Immuno-isolation of autophagic compartments was adapted from (14). Seedlings were homogenized in vesicle isolation buffer and subjected to differential centrifugation followed by sucrose cushion ultracentrifugation to isolate a membrane fraction (MF). Immuno-isolation was performed in native conditions using GFP-Trap beads targeting GFP-ATG8A or YFP-ATG18A (IP_+_). Control immunoprecipitations (IP_−_) were performed using uncoated beads. Isolated fractions were washed and processed for downstream proteomic and lipidomic analyses. Full experimental details are provided in **Supplemental Information**.

### Quality control of immuno-isolated fractions

The enrichment and purity of autophagic compartments were assessed by western blot using anti-GFP and anti-NBR1 antibodies and a set of organelle-specific markers (ER, mitochondria, nucleus, chloroplast, plasma membrane and tonoplast). In addition, the morphology of isolated structures was analyzed by transmission electron microscopy following immunogold labeling using anti-GFP antibodies. Full experimental details are provided in **Supplemental Information**.

### Proteomic analyses

Proteins from the input membrane fraction (MF) and immuno-isolated membranes were separated by SDS-PAGE, in-gel digested with trypsin and analyzed by nanoLC-MS/MS using an Orbitrap mass spectrometer. Raw data were processed using Proteome Discoverer and searched against the *Arabidopsis thaliana* protein database. Label-free quantification was performed to determine relative protein abundances. Proteins enriched in autophagic compartments were identified based on statistical comparisons of IP_+_/IP_−_ and IP_+_/MF ratios using t-tests and defined enrichment thresholds. Functional annotation and Gene Ontology enrichment analyses were performed using the STRING database (https://string-db.or). More details are provided in **Supplemental Information**.

### Lipid analyses

Lipids from input MF and immuno-isolated membranes were extracted and analyzed using complementary GC-MS and LC-MS/MS approaches. Fatty acids and sterols were analyzed following transmethylation and derivatization. Glycerolipids, sphingolipids and phosphoinositides were quantified by LC-MS/MS using appropriate internal standards. For all lipid classes, the composition of autophagic compartments was determined by subtracting values obtained from control immunoprecipitations (IP_−_) from those obtained in specific fractions (IP_+_). For PG quantification in roots and leaves of *ami:pgps1,2* mutant, details are provided in **Supplemental Information**.

### Transient expression and confocal microscopy

Candidate proteins were transiently expressed as GFP fusions in *Nicotiana benthamiana* leaves together with RFP-ATG8E using *Agrobacterium tumefaciens*-mediated transformation. After expression and autophagy induction, samples were imaged using a Zeiss LSM 880 confocal microscope. Colocalization analyses were performed using Fiji software. More information is provided in **Supplemental Information**.

### Statistical analyses

Statistical analyses were performed using GraphPad Prism 11.0.2 (GraphPad Software, La Jolla, CA, USA) with tests indicated in the figure legends. P-values are as follows: p-value>0.05 (non-significant, ns), *p<0.05, **p<0.01, ***p<0.005, and ****p<0.001.

## Acknowledgments

We thank Professor L. Jiang (The Chinese University of Hong Kong) for YFP-ATG18A seeds and Professor J. Joubès (University of Bordeaux) for helpful discussions.

